# Di-HAMP domains of a cytoplasmic chemoreceptor modulate nucleoid array formation and downstream signalling

**DOI:** 10.1101/2024.09.02.610768

**Authors:** P J Jazleena, Apurba Das, Annick Guiseppi, Fabian Debard, Jaya Sharma, Mutum Yaikhomba, Tâm Mignot, Emilia Mauriello, Pananghat Gayathri

## Abstract

In bacterial chemosensing, environmental cues are typically sensed by bacterial transmembrane receptors known as methyl accepting chemotaxis proteins (MCPs). MCPs form highly organized arrays using the bacterial membrane as a scaffold, which amplify the signals and transduce them into a cellular response. The FrzCD cytoplasmic receptor from *Myxococcus xanthus* is unique due to its ability to bind DNA and use the nucleoid as a scaffold to form arrays. In this study, we identified two HAMP (***<u>h</u>***istidine kinase, ***<u>a</u>***denylyl cyclase, ***<u>M</u>***CP and ***<u>p</u>***hosphatase) domains located between the DNA binding and signaling domains of FrzCD. *In vitro* experiments demonstrate that these HAMP domains restrict FrzCD to a dimeric form in solution and modulate FrzCD’s affinity for DNA, whereas the signaling domain stabilizes higher-order oligomeric assemblies upon DNA binding. Through fluorescence microscopy and analyses on *Myxococcus* social behavior, we demonstrate that the impact of FrzCD HAMP domains on DNA binding and oligomerization significantly influences group motility and development. Our results suggest that HAMP domains might have roles not only in signal transduction but also in the plasticity of chemosensory arrays. These observations illustrate mechanisms of regulation of a DNA-bound cytoplasmic array formed by a diffusible MCP.

**Significance Statement:** Our study identifies the presence of tandem HAMP domains in a cytoplasmic chemoreceptor, FrzCD, from *Myxococcus xanthus*, and highlights their role in dynamic receptor oligomerization on a DNA scaffold. By controlling receptor oligomerization and subsequently the array formation on the nucleoid, the tandem HAMP domains impart plasticity to the receptor arrays. Such plasticity governs cellular responses to external signals and dictate bacterial social behavior in group motility and multicellular structure formation.

## Introduction

Bacteria sense external signals and convey them into various cellular behaviors through intricate regulatory pathways known as chemosensory systems (CSS). Chemosensory systems regulate diverse behavioral responses, including taxis, gene expression, and development (1–3). The chemical signal is sensed by specialized chemoreceptors, referred to as ***<u>m</u>***ethyl-accepting ***<u>c</u>***hemotaxis ***<u>p</u>***roteins (MCPs). A canonical MCP comprises a periplasmic ligand-binding domain, followed by a transmembrane region, a HAMP (***<u>h</u>***istidine kinase, ***<u>a</u>***denylyl cyclase, ***<u>M</u>***CP and ***<u>p</u>***hosphatase) domain and a methyl-accepting (MA) signaling domain in its C-terminal cytoplasmic region (2). The HAMP domain is a homodimeric four-helix bundle (4), where each monomer comprises two antiparallel amphipathic helices, AS1 and AS2, joined by a ∼14 residue flexible loop as a connector (4–6). Structural studies revealed two different HAMP conformations with different helical registers, rotation and crossing angles that dictate opposite downstream signals in bacterial chemotaxis (7). The helices in the HAMP domain undergo relative movements when the MCP is bound to its ligand, as observed from the structural states of the NarQ receptor captured with and without the ligand nitrate (8).

A common feature of MCPs is their ability to oligomerize and form highly ordered hexagonal arrays, where each core complex is composed of two MCP trimer of dimers, two CheW docking proteins and one CheA dimer (9–11). In these arrays, MCP hexagons, consisting of six MCP trimers of dimers, are networked by CheA-CheW rings essential for the integrity of the system (12, 13). Receptor arrays are essential for the amplification of the initial signal, which is a direct consequence of the cooperative interactions between clustered chemoreceptors (12, 14, 15). While arrays have been described as universal among prokaryotic and archaeal chemoreceptors (16), their subcellular localization and distribution vary among different bacterial species. Membrane chemosensory arrays are the most common and can be visualized by fluorescence microscopy as clusters located at the cell poles. However, there are MCPs that lack the transmembrane domain and are located in the cytoplasm (17). FrzCD, the MCP of the Frz system of the gliding bacterium *Myxococcus xanthus*, is an example of cytoplasmic MCP (18, 19).

*M. xanthus* is a bacterium with a multi-stage developmental cycle that involves swarming, predation and aggregation into specialized multicellular biofilms termed fruiting bodies. Out of eight CSS and a total of 21 chemoreceptors in *M. xanthus* (20), the most studied CSS is the Frz system, which modulates the frequency at which cells reverse their direction of movement in a chemotactic-like manner (21). This regulation is critical for a successful developmental cycle because the aggregation of *M. xanthus* cells into fruiting bodies is dependent on the frequency of reversals (22). In addition to FrzCD, the cytoplasmic MCP, the Frz system comprises FrzE (analogous to CheA), FrzA and FrzB (analogous to CheW), two CheY-like response regulators, FrzX and FrzZ, as well as FrzF and FrzG (analogous to CheR and CheB, respectively) (22, 23). How environmental signals are perceived by FrzCD remains elusive as this protein lacks an obvious sensing domain.

Besides a conserved C-terminal methyl-accepting (MA) domain (residues 137 to 417), FrzCD features a N-terminal positively charged amino acid sequence (residues 1 to 30) responsible for DNA-binding (DB domain) (Supplementary Figure S1A) (19). The functions and fold of residues 31 to 136 are unknown. We have previously shown that FrzCD localizes into multiple clusters distributed along the nucleoid (18, 19, 24). Colocalization of FrzCD with the nucleoid is due to the direct interaction of the FrzCD N-terminal domain and DNA, in a DNA sequence-independent manner (19). Removal of the N-terminal region of FrzCD causes the loss of association with the nucleoid and the dispersal of FrzCD in the cytoplasm, indicating that the FrzCD binding to DNA is required for cluster formation (19). We also showed that the outcome of this nucleoid-mediated clustering of FrzCD is the same as that of transmembrane chemosensory arrays, which is the ability of the system to respond cooperatively to external signals (19).

In this study, we show the existence of two tandem HAMP domain units between the DNA-binding and the methyl-accepting coiled coil domains of FrzCD. We also show that the absence of one or both HAMP domains results in distinct modifications of Frz clusters on DNA in single cells; in altered reversal frequencies and, as a consequence, in motility and developmental defects. Our complementary *in vitro* data suggest that while the MA domain allows FrzCD oligomerization, the tandem HAMP domain restricts the oligomeric status of FrzCD in solution to a dimer. This implies that the tandem HAMP domains are essential for regulating the oligomerization status of the cytoplasmic FrzCD to a dimer until cluster formation occurs on the DNA. Thus, HAMP domains play a crucial role in regulating the binding of FrzCD to DNA, the exposure of eventually methylated residues, and thereby reversal frequencies. In summary, this study provides insights into the array formation of a cytoplasmic MCP, very few of which are characterized to date, and how DNA binding and array formation are modulated by the protein domain architecture.

## Results

### Structure prediction and homology show the presence of two tandem HAMP domains in FrzCD

Secondary structure predictions indicated that the N-terminal portion from residue 31 to 136 of FrzCD, forms alpha helices interrupted by two non-helical gaps (residues 49-61 and 106-118) (Supplementary Figure S1B, C). This observation led us to examine the sequence for the presence of multiple HAMP domains. While common domain prediction servers, such as Superfamily, CD-VIST or SMART (25–27) could not identify HAMP domains in FrzCD, a visual inspection, followed by sequence alignment with other poly-HAMP domains, revealed two consecutive HAMP domains, labeled as H1 (residue 31 to 83) and H2 (residue 84 to 137), with the AS2 helix of H1 in tandem with AS1 of H2 (Supplementary Figure S1B, C). The alignment also showed the presence of conserved glycine residues flanking the loop region (Supplementary Figure S1B).

A homology model of FrzCD showed a cylindrical structure with a length of approximately 300 Å (Figure 1A) and a diameter of approximately 14 Å (Figure 1B). The loops connecting the AS1 and AS2 helices of each HAMP domain extended outward from the cylinders formed by the helices (Figure 1B). The presence of two concatenated HAMP domains resulted in loops connecting AS1 and AS2 of the HAMP domains extending out of the FrzCD surface at varying angles, depending on the helix orientation. Structure predicted by AlphaFold2 aligned well with our homology model for FrzCD (Supplementary Figure S2A, B). The predominantly alpha helical content was validated by circular dichroism spectroscopy (Supplementary Figure S2C).

**Figure 1.**
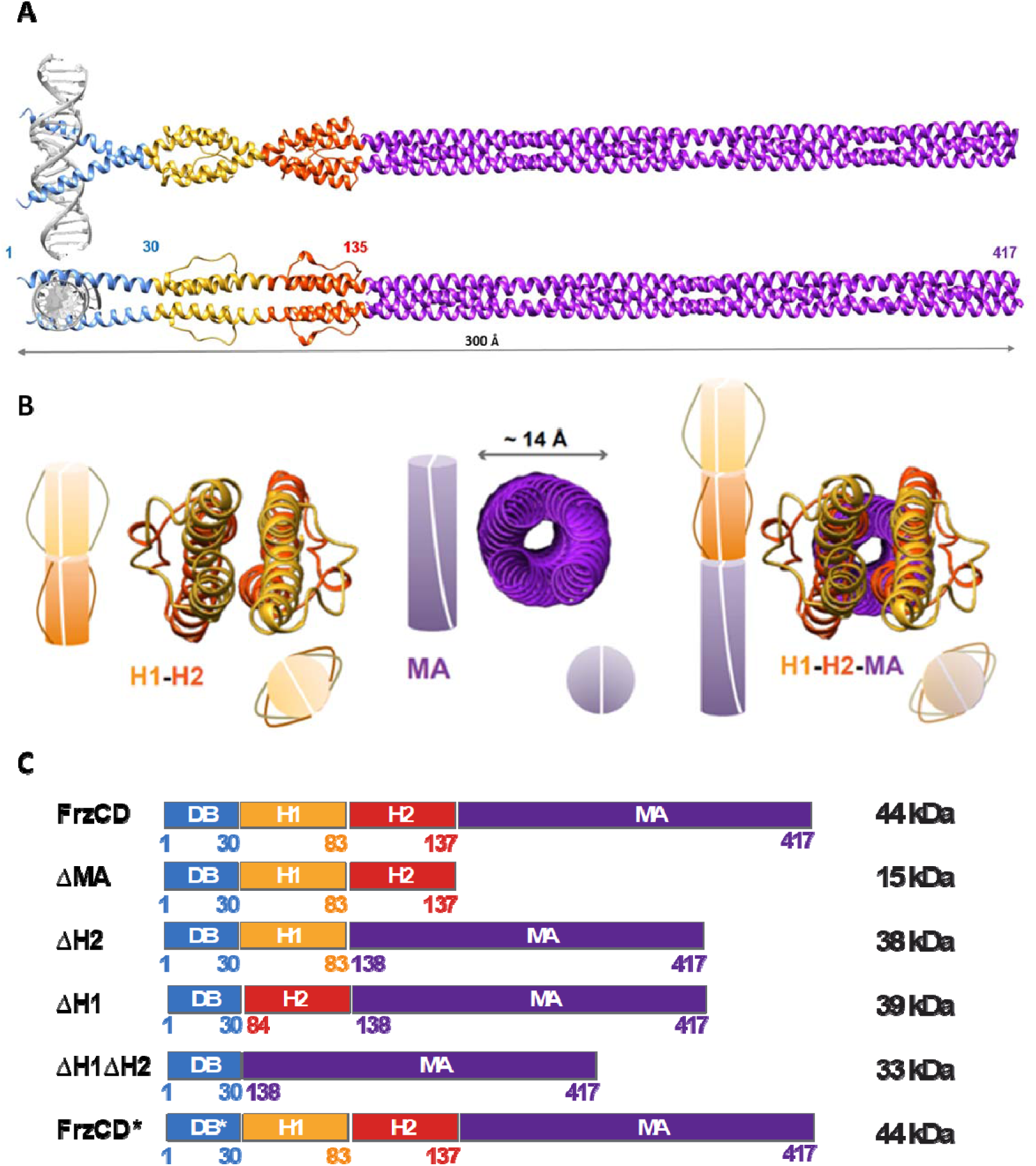
FrzCD possesses two HAMP domains between the DB and MA domains. **A.** FrzCD dimer model is an elongated structure of about 300 Å, generated as illustrated in Supplementary Figure S2B. Domain architecture of FrzCD comprises N-terminal DNA binding region (DB: residues 1 to 30) followed by a concatenated di-HAMP unit composed of HAMP1 (H1: residues 31 to 83) and HAMP2 (H2: residues 84 to 137) terminating in a long C-terminal coiled-coil signaling domain (MA: 40H, residues 138 to 417). **B.** Schematic representation of side and end-on views of the dimeric FrzCD domains. An end-on view of the dimeric model shows that the loops connecting the AS1 and AS2 helices of H1 and H2 project out from the cylindrical surface formed by the helices. The MA domain subtends a circular view of approximately 14-Å diameter. The loops connecting AS1 and AS2 of the two HAMP domains may extend out onto the surface at varying angles depending on the relative orientation of the helices. **C.** Schematic representation of various FrzCD domain deletion and mutant constructs and their corresponding molecular weights. DB stands for DNA-binding domain, H1 and H2 denotes HAMP domains, and MA for methyl-accepting domain. Numbers indicate the amino acid positions.

These results allowed us to redefine the FrzCD domain architecture, which included a newly identified tandem di-HAMP unit comprising H1 (residues 31 to 83) and H2 (residues 84 to137), positioned between the DB (bZIP-like) and the MA domains (Supplementary Figure S2D, E). We proceeded to validate our model experimentally using domain-wise deletion constructs (Figure 1C).

### HAMP domains of FrzCD modulate the methylation-mediated regulation of reversal frequency

To probe the role of FrzCD HAMP domains in protein activity and protein localization at the nucleoid *in vivo*, we first constructed a FrzCD-mNeongreen (FrzCDmNG) fusion. Similar to a previous FrzCD-GFP construct ((18–20, 24), FrzCDmNG exhibited stable expression and formed clusters at the nucleoid when expressed at the endogenous locus and under the *frz* promoter, but was not fully functional in mediating swarming (Supplementary Figure S3A-C). However, when the same fusion was expressed under a promoter inducible by vanillate at a heterologous locus (*Pvan-frzCDmNG*), it fully complemented the absence of the native copy of *frzCD* (Supplementary Figure S3C). The compromised functionality of FrzCDmNG when expressed at the endogenous locus was due to a 7.84 ± 0.96 log_10_-fold increase in expression (as measured by q-RT PCR) of the co-transcribed downstream gene, *frzE*, compared to the wildtype. Finally, the ability of *P_van_*-*frzCDmNG* to form clusters was indistinguishable from that observed for *frzCDmNG* at the endogenous locus (Supplementary Figure S3B).

Next, we systematically deleted either or both of the HAMP domains from the *P_van_-frzCDmNG* construct (Figure 1C) and induced the various alleles in a Δ*frzCD* strain. The deletion of either H1 or H2 resulted in distinct phenotypes, indicating that these two domains are functionally non-redundant. Single cell motility behaviors suggest that the motility defect observed in the absence of H1 is probably attributable to decreased single-cell reversal frequencies measured to be as low as those of Δ*frzCD* cells (Figure 2A-B). On the other hand, the motility defects of cells lacking H2 were due to increased reversal frequencies, reaching approximately twice that of wild type (Figure 2A-B). Additionally, while reduced reversals did not affect fruiting body formation in *P_van_-frzCDmNG*^Δ*H1*^, increased reversals prevented the formation of fruiting bodies in *P_van_-frzCDmNG*^Δ*H2*^, similar to other strains with increased reversals (Figure 2A-B) (22). *P_van_-frzCDmNG*^Δ*H1*Δ*H2*^ phenocopied *P_van_-frzCDmNG*^Δ*H1*^ in reversal frequencies and fruiting body formation (Figure 2A-B).

**Figure 2.**
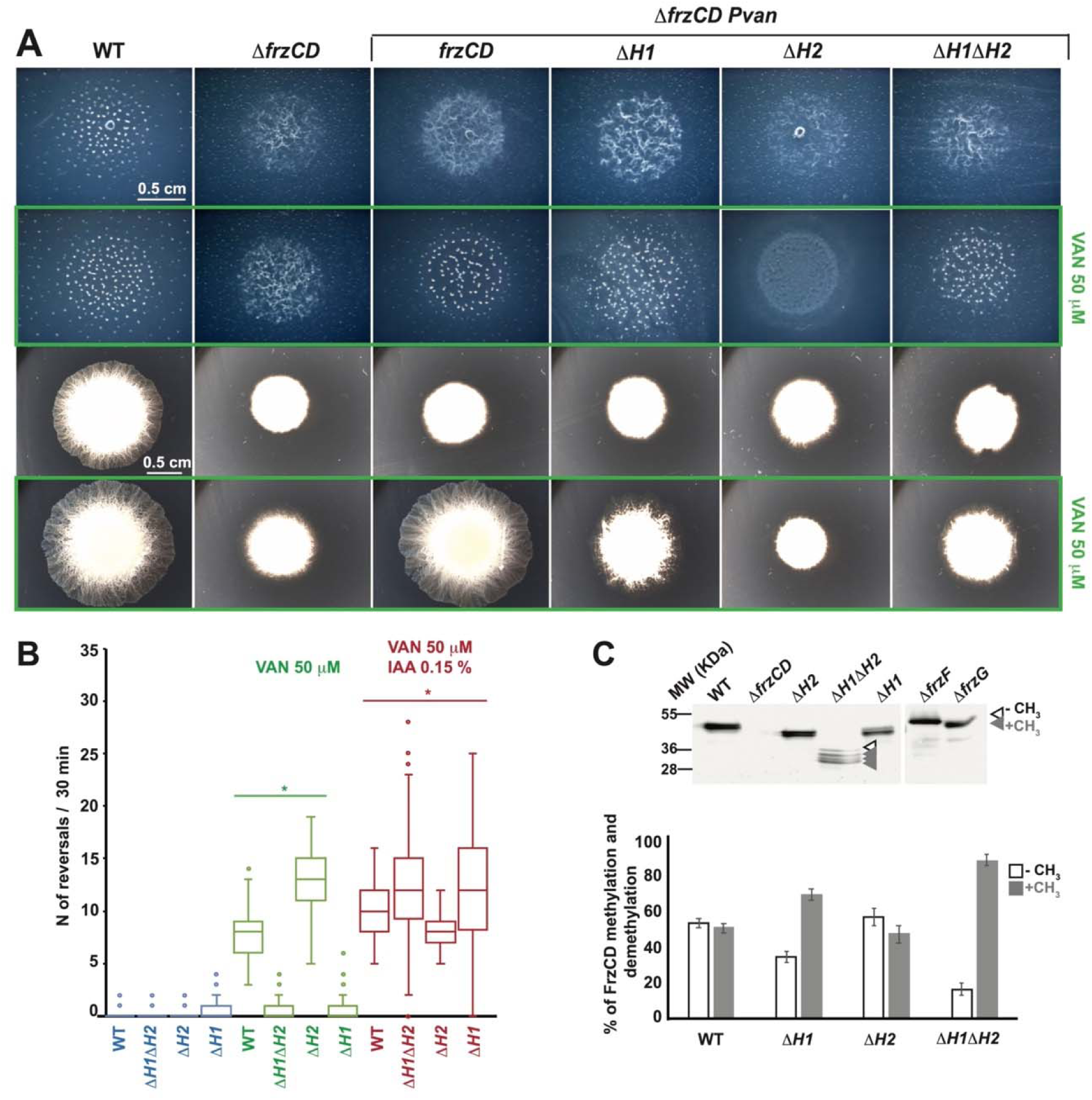
FrzCD H1 and H2 domains are important in the methylation-mediated regulation of reversal frequency and motility. **A.** Representative phenotypic assays of DZ2 (wild type), EM525 (Δ*frzCD*), EM885 (Δ*frzCD pvanfrzCDmNG*), EM914 (Δ*frzCD pvan-frzCDmNG*^Δ*H1*^), EM913 (Δ*frzCD pvan-frzCDmNG*^Δ*H2*^) and EM911 (Δ*frzCD pvan-frzCDmNG*^Δ*H1*Δ*H2*^). The first two rows are fruiting body formation assays on 1.5% CF agar, in the presence (green rectangle) or not of 50 µM vanillate, imaged at 72h. The last two rows are motility phenotypes on 0.5% CYE agar, in the presence (green rectangle) or not of 50 µM vanillate, imaged at 48h. **B.** Box plots of reversal frequencies of EM885 (Δ*frzCD pvan-frzCDmNG*), EM914 (Δ*frzCD pvan-frzCDmNG*^Δ*H1*^), EM913 (Δ*frzCD pvan-frzCDmNG*^Δ*H2*^) or EM911 (Δ*frzCD pvan-frzCDmNG*^Δ*H1*Δ*H2*^) single cells moving on agar pads supplemented or not with 50 µM vanillate or 50 µM vanillate and 0.15% IAA. The lower and upper boundaries of the boxes correspond to 25^th^ and 75^th^ percentiles, respectively. The median is shown as a line at the center of each box, and rounds represents the 10^th^ and 90^th^ percentiles. For the reversal frequency measurements, 100 cells from two biological replicates were used. **C.** Representative western blot on whole cell extracts of strains DZ2 (wild type), EM525 (Δ*frzCD*), EM775 (*frzCD*^Δ*H2*^), EM776 (*frzCD*^Δ*H1*Δ*H2*^), EM889 (*frzCD*^Δ*H1*^), EM527 (Δ*frzF*) and EM528 (Δ*frzG*), showing the presence of bands corresponding to methylated (grey arrowheads) and demethylated (white arrowheads) forms of FrzCD. The histogram shows the average percentage of intensity of methylated and demethylated bands on the total FrzCD intensity amount. Results are issued from three independent biological replicates. The error bars correspond to standard deviations.

The ability of cells to modulate their reversal frequencies is contingent on the methylation state of FrzCD, wherein low methylation levels correlate with high reversal frequencies and, conversely, high methylation with fewer reversals (28). Therefore, we sought to check whether the reversal frequency defects of our FrzCD HAMP mutants were also correlated with their methylation states. Given the difficulties in resolving methylation in FrzCDmNG by western blot due to its high molecular weight, we generated in-frame deletions of each or both HAMP domains at the endogenous locus and performed methylation assays. In these experiments, the highest band corresponds to the unmethylated state, while all other bands to different degrees of methylation with the lowest band corresponding to the fully methylated state (29, 30). In the case of FrzCD^ΔH1^, methylation levels were higher than wild type, while for FrzCD^ΔH2^ demethylation was prevalent (Figure 2C). Furthermore, for FrzCD^ΔH1ΔH2^ the methylation levels were similar to FrzCD^ΔH1^. These results showed that the absence of the HAMP domains altered the methylation of FrzCD, with further downstream effects.

The simultaneous deletion of H1 and H2 resulted in phenotypes similar to those of *Pvan-frzCDmNG*^Δ*H1*^, implying a dominant effect of Δ*H1* on Δ*H2* (Figure 2A-C). Notably, *Pvan-frzCDmNG*^Δ*H1*^, *Pvan-frzCDmNG*^Δ*H2*^ and *Pvan-frzCDmNG*^Δ*H1*Δ*H2*^ single cells all responded to a known activator of FrzCD, isoamyl alcohol (IAA), by frequently reversing their movement direction, similarly to wild type (Figure 2B). This suggests that all the above protein variants were able to signal to downstream Frz proteins in the presence of high signal levels. The phenotypes of HAMP mutants were not due to expression defects of the *frzCD* alleles (Supplementary Figure S3D). Together, these results suggested that the absence of the HAMP domains altered FrzCD activity, but in the presence of high signal concentrations, the ability of the FrzCD variants to transduce signals remained unchanged compared to the wild type.

### FrzCD HAMP domain deletions have differential effects on cluster formation upon nucleoid binding

To establish a correlation between the role of HAMP domains in FrzCD activity (Figure 2) and FrzCD localization to the bacterial nucleoid, we looked at the cellular localization of different FrzCD HAMP mutants fused to the Neongreen. Similar to the previously described FrzCD-GFP (19), FrzCD-mNG formed multiple clusters that were evenly distributed on the nucleoid (Figure 3 and Supplementary Figure 4A). Deletions of each or both HAMP domains did not impede either cluster formation or binding to DNA, as *M. xanthus* cells bearing these deletions all formed Frz clusters at the nucleoid (Figure 3 and Supplementary Figure 4A). However, the functions of H1 and H2 in cluster formation were not redundant, as cells with either deletion exhibited clusters with different localization patterns. In the absence of H1, clusters resembled those formed by FrzCD-mNG in intensity and distribution (Figure 3 and Supplementary Figure S4), although they appeared less defined (Figure 3C). In the absence of H2, cells formed fewer clusters but with high fluorescence intensities (Figure 3 and Supplementary Figure S4). Finally, the deletion of both HAMP domains resulted in the formation of highly condensed clusters (Figure 3), slightly more numerous than wild type (Supplementary Figure S4). The fluorescence intensity of FrzCD^ΔH1ΔH2^-mNG clusters was similar to FrzCD-mNG, but cells showed less diffused fluorescence, suggesting that all FrzCD^ΔH1ΔH2^-mNG protein units are engaged in clusters (Figure 3).

**Figure 3.**
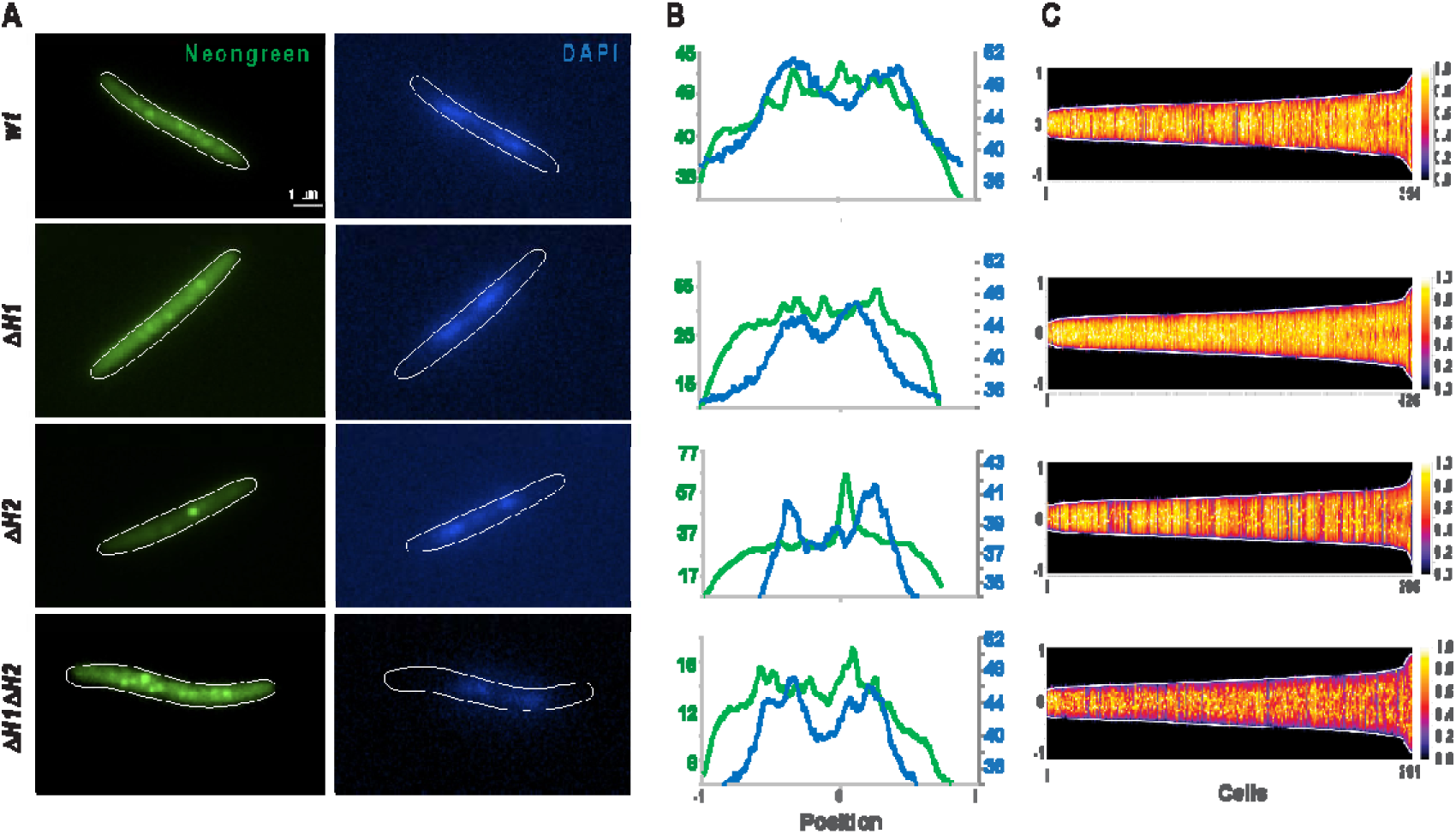
The HAMP domains play a role in FrzCD cluster formation on the nucleoid. **A.** Representative micrographs of *M. xanthus* cells from strains EM885 (Δ*frzCD pvan-frzCDmNG*), EM914 (Δ*frzCD pvan-frzCDmNG*^Δ*H1*^), EM913 (Δ*frzCD pvan-frzCDmNG*^Δ*H2*^) or EM911 (Δ*frzCD pvan-frzCDmNG*^Δ*H1*Δ*H2*^) stained with Neongreen and the DNA DAPI stain. The genetic backgrounds of the *M. xanthus* strains are indicated on the left. Scale bars correspond to 1 μm. **B.** Neongreen (green) and DAPI (blue) fluorescence profiles with the fluorescence intensity (arbitrary units) represented on the y axis and the cell length positions with -1 and +1 indicating the poles and 0 the center, on the x axis. **C.** For each indicated strain, at least 290 cells (x axis) from at least two biological replicates are represented as lines and ordered according to their length (pixels) in demographs. The Neongreen fluorescence intensity along the cell body is represented as colored pixels at the corresponding cell position (from -1 to +1 on the y axis). “0” is the cell center. On the right, a scale indicates the fluorescence intensity and the corresponding colors.

In conclusion, HAMP domains were not necessary for cluster formation, but their absence appeared to impact cluster intensity and number.

### HAMP domains reduce the affinity of FrzCD for DNA, the MA domain favors FrzCD-DNA stable complexes

The observed changes in cluster formation among the various HAMP mutants could be due to impaired DNA-binding. To estimate the affinity of FrzCD for DNA in the presence or absence of its HAMP domains, we heterologously expressed and purified various His_6_-tagged FrzCD domains, along with the full-length protein. The purification yielded stable and soluble proteins, supporting a reliable prediction of domain boundaries (Supplementary Figure S5).

As we have shown previously, the interaction of FrzCD with DNA is charge-dependent and independent on DNA sequence, length (from 69 to 1,254 bp) and GC content (19). We compared the binding of HAMP deletion constructs of FrzCD with the wild type. By performing electromobility shift assays (EMSA), we confirmed that full-length FrzCD bound to DNA fragments as short as 8 bp and up to 600 bp (Supplementary Figure S6, S7A). FrzCD could also bind to 8-base single-stranded DNA fragments (Supplementary Figure S7B), strengthening the hypothesis that the FrzCD-DNA interaction is predominantly charge-based and involves the DNA phosphate backbone. To provide direct evidence for this charge-mediated interaction, we replaced three lysine (K9, K13 and K18) and two arginine (R15 and R17) residues of FrzCD by the negatively-charged glutamates (FrzCD*). The DNA binding capacity of this FrzCD* variant was completely abolished (Supplementary Figure S7C), confirming that the FrzCD-DNA interaction is predominantly charge-mediated.

FrzCD^ΔMA^ exhibited stable binding comparable to the wild type, only to very short DNA fragments of 8 bp to 12 bp (Supplementary Figure S6). Smearing was observed for DNA fragments longer than 12 bp, indicative of unstable complex formation. While 8 bp is probably the minimal footprint of a dimeric helical structure (31), larger DNA fragments might require a stable FrzCD oligomerization for the DNA-FrzCD complex formation. The MA domain might thus be important for the stabilization of FrzCD oligomers on large DNA fragments (Supplementary Figure S6, S7A). Comparison with the binding profile of FrzCD* variant (Supplementary Figure S7C) indeed demonstrated that the smears observed in FrzCD^ΔMA^ did not indicate a complete abrogation of DNA binding, but an unstable complex.

Next, we compared the affinity of FrzCD for DNA in the presence or absence of its HAMP domains. Stable shifts were observed for FrzCD constructs lacking one or both HAMP domains (FrzCD^ΔH2^, FrzCD^ΔH1^ or FrzCD^ΔH1ΔH2^) with DNA fragments from 35 to 432 bp (Figure 4A). This supported the *in vivo* observation that all these FrzCD variants mediated cluster formation on the nucleoid (Figure 3). Interestingly, the absence of both HAMP domains significantly improved the affinity for DNA as compared to wildtype and single HAMP deletions (Figure 4A).

**Figure 4:**
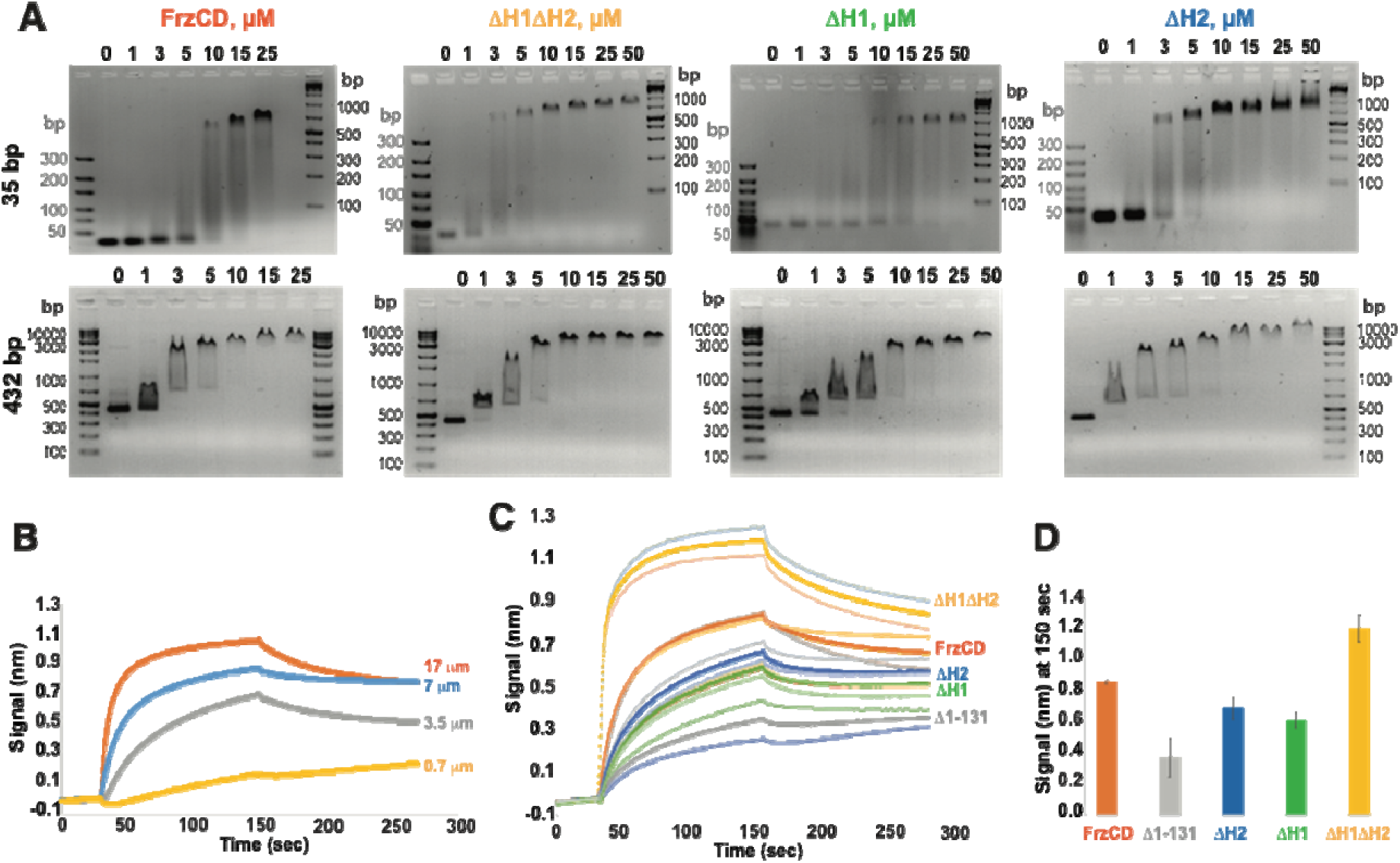
HAMP domains reduce the affinity of FrzCD for DNA. **A.** Representative electrophoretic mobility shift assays (EMSA) resolving the binding to DNA. The indicated concentrations of purified FrzCD protein domains as well as FrzCD full-length were incubated with 0.5 µM DNA fragments of the indicated sizes. DNA ladders are indicated as base pairs. **B.** Average binding curves of increasing concentration of 6His-FrzCD with an immobilized 178 bp DNA fragment at a concentration of 5 nM. **C.** Average binding curves and biological replicates in color gradations of the indicated His_6_-FrzCD variants at 7 mM concentration, with an immobilized 178 bp DNA fragment at a concentration of 5 nM. **D.** Average binding signals at 150 sec, extrapolated from (C) for the indicated FrzCD variants. Error bars indicates standard deviations for two biological replicates. The sequences of the DNA fragments used in panels A-D is indicated in Supplementary Table S4.

To quantify the FrzCD-DNA binding and explore the dynamics of the FrzCD binding to DNA, we used Bio-layer Interferometry (BLI), as previously described (19, 24).The FrzCD construct lacking its N terminus (FrzCD^Δ1-131^; MA domain only) served as negative control. First, we optimized the binding of FrzCD to the biosensor by testing different FrzCD concentrations (Figure 4B). Using this concentration range, we determined the K_D_ of FrzCD for DNA to be 1.074 μM. Then, we tested the DNA-binding of FrzCD mutants lacking the HAMP domains. In line with the gel-shift experiments, FrzCD^ΔH1ΔH2^ showed a higher affinity for DNA than the wild type (0.203 μM vs 1.074 μM) (Figure 4C-D). FrzCD^ΔH1^ and FrzCD^ΔH2^ exhibited slightly decreased affinity for DNA compared to wildtype (Figure 4C-D), with K_D_ values of 2.27 μM and 2.64 μM, respectively. Finally, as expected, no DNA binding was detected when FrzCD^Δ1-131^ was used (Figure 4C-D).

### Tandem HAMP domains restricts oligomerization in the absence of DNA

The notion that transmembrane MCPs form high-ordered structures, coupled with the observation of FrzCD-DNA complexes of increasing molecular weights (supershifts) in EMSA, suggest the formation of FrzCD oligomers on DNA. Moreover, supershifts are also evident in the absence of the MA and HAMP domains (Figure 4 and Supplementary Figure S6, S7A). Hence, we decided to test the oligomerization of FrzCD constructs in the absence and presence of the DNA scaffold. Given that FrzCD has an elongated shape and is not globular, we employed size exclusion chromatography (SEC) combined with multi-angle light scattering (SEC-MALS) measurements to accurately estimate the oligomeric status. Results indicate that full-length FrzCD and FrzCD^ΔMA^ exist only as dimers even at input concentrations as high as 150 and 200 µM, respectively (Figure 5A). Conversely, FrzCD^ΔH1^, FrzCD^ΔH2^ or FrzCD^ΔH1ΔH2^ form higher oligomers, showing the presence of tetramers even at lower protein concentration (Figure 5A). These observations suggest that HAMP domains prevent the formation of FrzCD oligomers *in vitro*.

**Figure 5:**
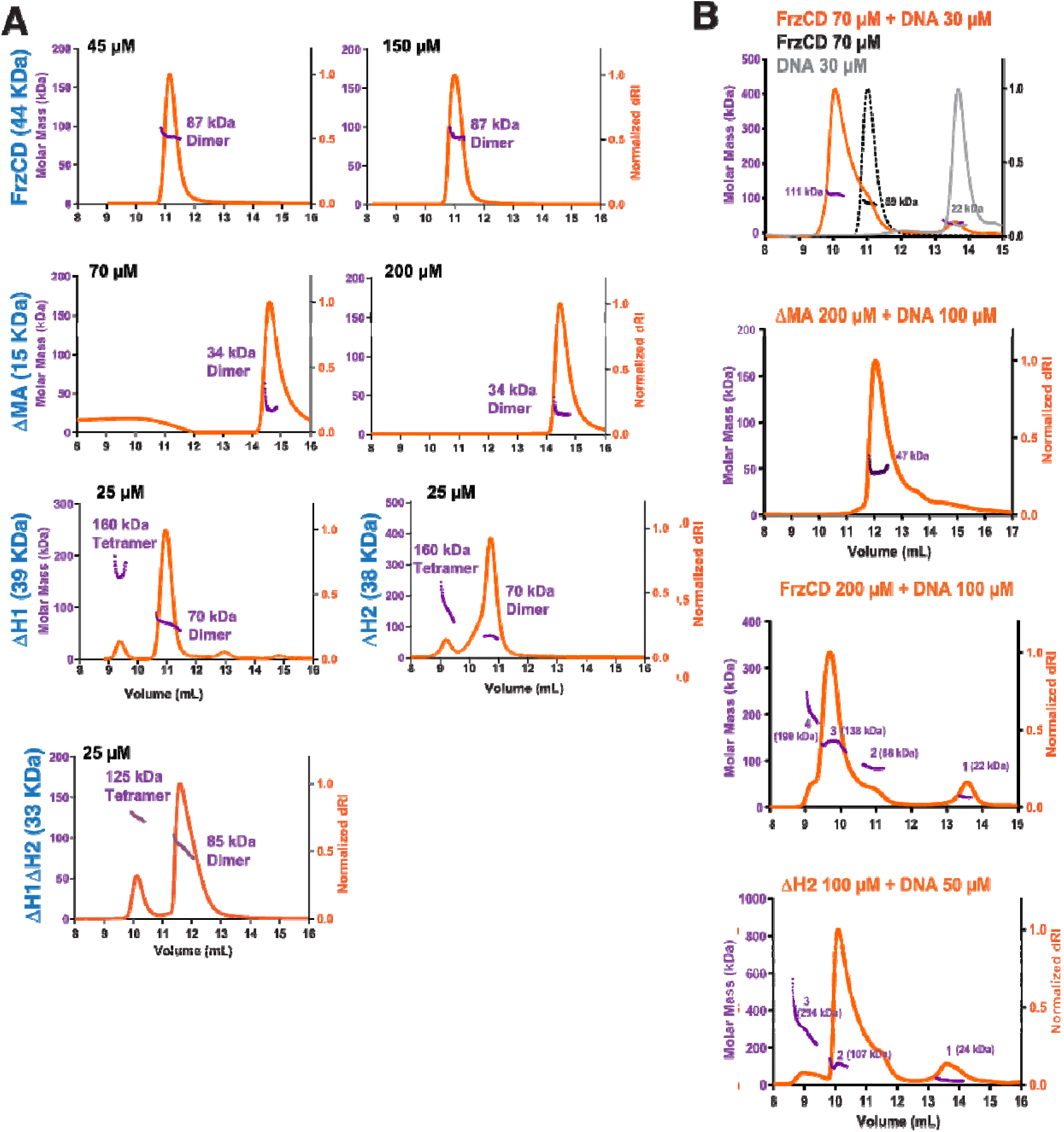
HAMP domains prevent FrzCD oligomer formation in the absence of DNA. **A.** SEC-MALS profiles for the indicated forms of FrzCD. Proteins were used at the indicated concentrations. The axes read as in **B.** **B.** SEC-MALS profile for the indicated forms of FrzCD in the presence of a 35-bp DNA. Proteins and DNA were used at the indicated concentrations. The y-axes represent molar mass (left, purple) and normalized differential refractive index (dRI, orange) while the x-axis denotes the volume of elution. The molecular masses corresponding to each peak are labeled. Supplementary Table S5 summarizes the calculated molecular weights based on the SEC-MALS experiments, and the expected molecular weights of the protein-DNA complexes.

To investigate whether the inability of FrzCD^ΔMA^ to form stable complexes with high molecular-weight DNA was due to an oligomerization defect, we decided to characterize the oligomerization status of both FrzCD and FrzCD^ΔMA^ upon DNA binding. For the SEC-MALS experiments, a 35-bp oligonucleotide was used so that the elution profiles of free DNA and free protein were well-resolved (Figure 5B). Upon addition of DNA, both FrzCD and FrzCD^ΔMA^ eluted as complexes with DNA, but, most strikingly, FrzCD formed high-order oligomers at high protein concentration (Figure 5B), unlike in the absence of DNA (Figure 5A). On the other hand, FrzCD^ΔMA^ could only be resolved as dimer, even in complex with DNA and at high protein concentrations (Figure 5B). This observation is consistent with the EMSA results, where supershifts were not observed for FrzCD^ΔMA^ (Supplementary Figure S6). Lastly, FrzCD^ΔH2^ also formed high-order oligomers in the presence of DNA (Figure 5B), behaving similarly to conditions in the absence of DNA (Figure 5A). Supplementary Table S5 summarizes the calculated molecular weights based on the SEC-MALS experiments, and the expected molecular weights of the protein-DNA complexes. Together, these results indicate that while the MA domain is required for FrzCD oligomerization, HAMP domains might prevent such oligomerization when DNA is absent.

## Discussion

MCP array formation is an essential feature for the allosteric activation of the chemosensory pathway and the generation of amplified responses for functioning over a wide range of signal concentrations (12). In the absence of a transmembrane domain, the cytoplasmic MCP FrzCD utilizes the nucleoid as a scaffold to form dense, dynamic clusters (19, 24, 32). The dynamic nature of these clusters likely stems from the intrinsic plasticity of FrzCD arrays (24), resulting from a delicate equilibrium between oligomerization and molecular turnover among arrays. In this work, we report the presence of two in-tandem HAMP domains connecting the DB and MA signaling domains of FrzCD. The existence of a HAMP domain in FrzCD has already been proposed in an earlier study (33). However, the exact boundaries of this putative HAMP domain, roughly aligning with our H2 domain, remained imprecise due to the absence of detailed structure predictions (33). Here, through a combination of structure predictions and careful visual sequence comparison, we identified two sequential HAMP domains, H1 and H2. These domains exhibit conserved non-helical loops that extend out of the elongated and predominantly helical structure of FrzCD.

We show that a presence of the tandem HAMP domains primarily prevents the formation of higher order oligomers of FrzCD in the absence of DNA, based on *in vitro* experiments. We also show that the DNA-dependent oligomerization beyond the dimeric state strictly depends on the presence of the MA domain. We, thus, propose a model where, following the initial DNA-binding primarily by the positively charged alpha helix at the FrzCD DB domain, conformational changes are transduced to the HAMP domains and, subsequently, to the MA domain. These conformational changes lead to differential exposure of the residues of the MA domain for methylation. Also, the MA domain can, in this new state, probably mediate the formation of trimers of dimers (Figure 6). Additionally, HAMP domains influence DNA binding capability of FrzCD. In fact, we demonstrate that the deletion of both H1 and H2 together significantly enhances the affinity of FrzCD for DNA *in vitro*. This behavior is likely linked to the formation of well-defined and intense, albeit non-functional nucleoid-clusters *in vivo* (Figure 6). Similarly, the reduced ability of FrzCD^ΔH1^ and FrzCD^ΔH2^ to form clusters on the nucleoid *in vivo*, with FrzCD^ΔH1^ clusters appearing more diffused compared to wildtype and FrzCD^ΔH2^ forming fewer clusters, may be attributed to their diminished DNA-binding capability.

**Figure 6:**
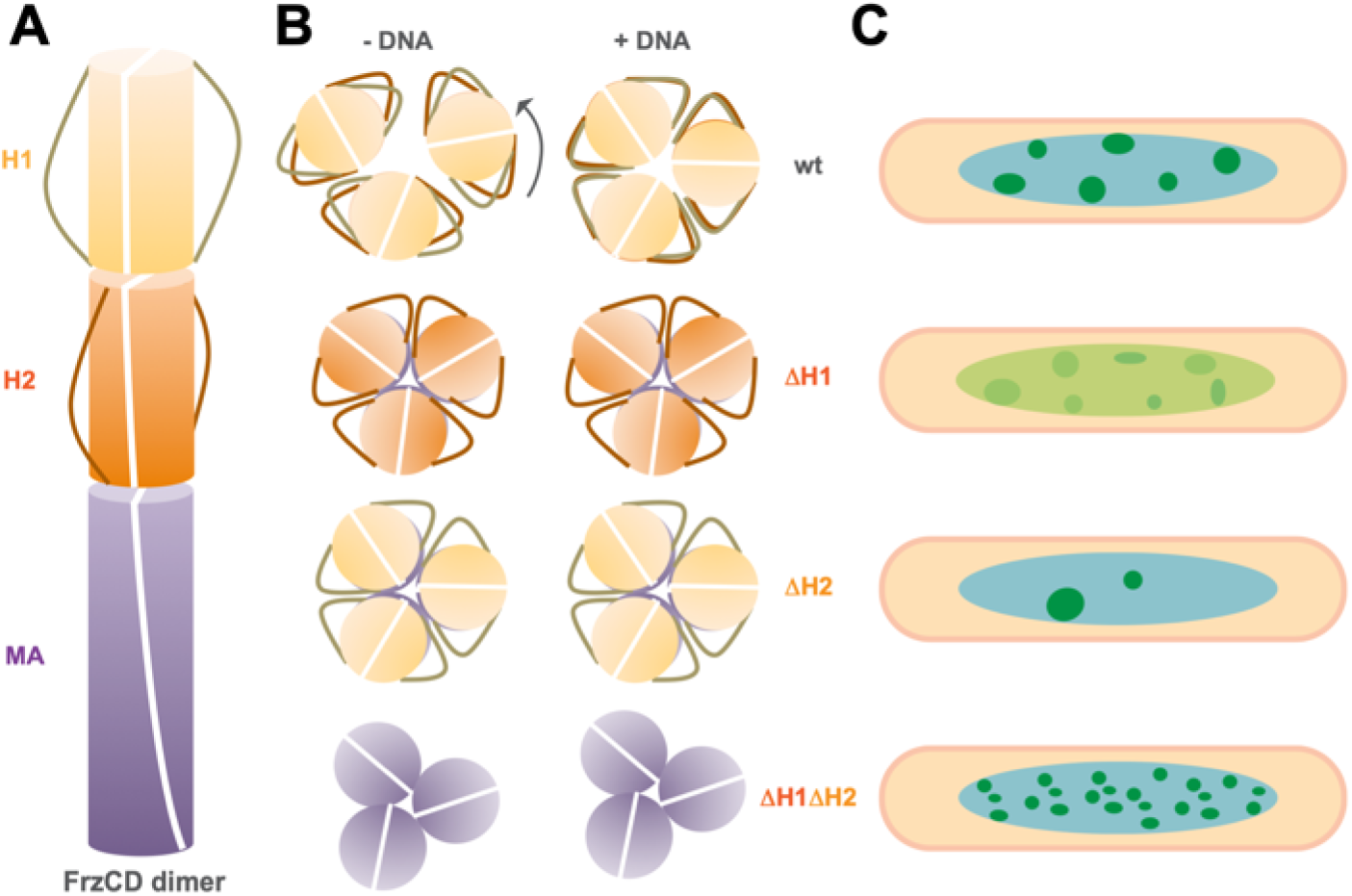
Model for the DNA-mediated regulation of FrzCD oligomerization. **A-B.** Schematic representation of FrzCD dimer and individual domains. **A.** side view, **B.** top view. **B.** In wildtype, the orientation of the loop regions of the tandem HAMP domains restricts the formation of trimers of dimers unless DNA binding causes rotation of the helices. The presence of a single or no HAMP domain allows the formation of trimers of dimers even in the absence of DNA. **C.** Altered FrzCD oligomerization and affinity for DNA in the absence of one or both HAMP domains have effects on FrzCD cluster formation on the nucleoid. When both H1 and H2 are deleted the affinity of FrzCD for DNA largely improves *in vitro*, likely explaining the formation of well-defined and bright, although non-functional nucleoid-clusters. Similarly, while the deletion of H1 or H2 allows oligomerization even in the absence of DNA, the FrzCD affinity for DNA is reduced in these mutants. This likely explains more diffused clusters in FrzCD^ΔH1^ and fewer clusters in FrzCD^ΔH2^ as compared to wildtype.

The precise molecular mechanisms through which HAMP domains restrict the oligomerization state of FrzCD to a dimer in solution, thereby preventing higher-order oligomerization in the absence of DNA, can be explained by considering the following model. As illustrated in Figure 6 and based on our predictions of the FrzCD dimer structure, the non-helical loops that connect AS1 and AS2 of H1 and H2 extend out of the elongated helical structure of FrzCD. This could hamper the formation of trimers of dimers, the canonical organization of chemoreceptors, when FrzCD is free in the cytoplasm. We hypothesize that when FrzCD encounters the nucleoid scaffold, DNA-binding induces a conformational change characterized by an axial rotation of the HAMP helices. This conformational change might alleviate the steric hindrance to oligomer formation imposed by the protruding loops. Additionally, this rotation may be transduced to the MA domain, allowing the exposure of an interface responsible for the formation of trimers of dimers (Figure 6). The interplay between DNA-binding events and consequent structural dynamics contributes to the regulation of FrzCD oligomerization states in response to its cellular environment.

The proposed mechanism is consistent with what has been reported for the molecular role of HAMP domain in chemoreceptor activation, where the rotation of AS1 and AS2 is responsible for transducing signals from the sensing to the MA domain (5). Our model is also coherent with what reported for poly HAMP domains, where molecular movements including helix rotation, translation or tilt, facilitate transitions between HAMP conformers (7). In the case of FrzCD, the tandem arrangement of HAMP domains might favor the optimal orientation of the protruding loops, effectively preventing oligomerization in the cytoplasm before contacts with DNA. This specific arrangement of HAMP domains in FrzCD might have evolved to ensure signal transduction when the membrane MCP scaffold is replaced by DNA. A better understanding of the sensing mechanisms of FrzCD in the future will likely contribute to further explain the requirement of two HAMP domains.

The rotation model we propose might also explain the methylation and phenotypic defects observed in the HAMP deletion strains. According to our model, the putative rotation of the HAMP domains, transduced to the MA domain, might influence the accessibility of the methylation sites on the MA domain. Impaired methylation states result, in turn, in alterations in reversal frequencies and social behaviors.

Two additional interesting results emerged from our *in vitro* study. First, the dispensability of the MA domain in the formation of FrzCD dimers implies that the DB and HAMP domains alone are sufficient for FrzCD dimerization. Interestingly, our EMSA assays show that both FrzCD and FrzCD^ΔMA^ can efficiently form complexes with DNA fragment as short as 8 bp. This DNA length corresponds to the minimal footprint of a dimeric helical structure (31), suggesting that the DB domain could bind DNA in a dimeric form. The second finding suggests that in the presence of MA domain, FrzCD can form oligomers *in vitro* even without its downstream Frz partners. However, *in vivo,* clusters of FrzCD were observed very rarely in the absence of FrzA and FrzE (19, 24). This suggests that other factors are likely required for efficient array formation and its stability *in vivo*, even in the presence of the MA domain.

In conclusion, this study brings new insights into the role of HAMP domains, not only in signal transduction but also in the plasticity of chemosensory arrays, establishing the delicate equilibrium between oligomerization and molecule turnover. The presence of two consecutive HAMP domains emerges as an intrinsic necessity, when scaffolds other than the inner membrane are utilized to nucleate receptor arrays. This introduces a novel dimension to the understanding of these universally conserved domains, revealing their potential significance beyond their conventional role in signal transduction.

## Materials and Methods

### Domain architecture prediction for FrzCD

Secondary structure of FrzCD was predicted using Psipred (34) and Jpred 4 (35). Further, servers such as Superfamily (25), CD-VIST (26) or SMART (27) were employed to determine the domains of FrzCD. A multiple sequence alignment (MSA) was performed for FrzCD against poly-HAMP domains of other MCPs, such as Aer2 and HAMP consensus sequences obtained from the work by Dunin-Horkawicz and Lupas (36) using Jalview software (Supplementary Figure 1B) (37). The alignment was inspected for presence of any HAMP domains. Based on the secondary structure prediction as well as MSA (Supplementary Figure 1C), domain boundaries were predicted for FrzCD.

### Modeling of FrzCD

FrzCD was dissected into the DNA-binding (DB), di-HAMP (H1 and H2) and methyl-accepting (MA) domains. The N-terminal DNA-binding domain sequence was aligned against various protein structures from basic leucine zipper (bZIP) families such as 1GU4, Creb1, ATF4, Fos, Jun using Jalview software (Supplementary Figure 2D). To obtain a homology model for FrzCD, the approach included modeling each of these domains separately as dimers with a 12-residue overlap to put the model together. Therefore, for DB, 1-42 residues were modeled against GCN4 (PDB: 2dgc) while the di-HAMP domain (residues 31-135) was modeled against HAMP1 and HAMP2 of Aer2 (PDB: 3lnr) using SWISS-MODEL (38). Four helix bundle of MA (residues 124-417) dimer was generated using CCBuilder (39). Dimers of the three domains were superimposed based on C-α overlap regions. Each domain was individually modeled with a 12-residue overlap (equivalent to three alpha-helical turns), and subsequently, superimposition using Cl7 atoms (Supplementary Figure 2E) allowing to obtain a model of the entire FrzCD dimer. The domains were joined at residues Ser34 and Ser35 between DB and H1, and residues Val130 and Ile131 between H2 and MA to generate the FrzCD homology model. Recently, dimeric model of FrzCD was also generated in AlphaFold2 using ColabFold (Supplementary Figure 2B) (40, 41).

### Bacterial strains and plasmids

The strains and plasmids used in this work are listed in Supplementary Table S1 and Table S2, respectively. To create *M. xanthus* in-frame deletion strains, 900 bp upstream and downstream of the region targeted for deletion were amplified by PCR and cloned into the pBJ114 vector (22) previously digested with HindIII and EcoRI. To fuse FrzCD to the Neongreen reporter, *neongreen* was amplified and cloned into pDPA20 (18) previously digested with NotI and XhoI, to replace *gfp*. To generate *M. xanthus* strains expressing *frzCD-neongreen* alleles at the endogenous locus, *frzCD* variants plus an upstream region for a total of 900 bp, *neongreen* and 900 bp downstream of *frzCD* on the *M. xanthus* chromosome, were amplified and cloned into pBJ114 previously digested with HindIII and EcoRI. To obtain the plasmids for the inducible expression, the different *frzCD-neongreen* alleles were amplified from each respective *M. xanthus* strain and cloned into pMR3690 (42) previously digested with NdeI and BglII. *M. xanthus* competent cells were prepared and transformed by electroporation as described previously (22). Cells were grown in CYE rich medium containing, when required, kanamycin at a final concentration of 25 µg/ml. Vanillate was used to induce the different *frzCD-neongreen* alleles at a concentration ranging from 0.5 to 50 µM.

*Escherichia coli* DH-5α (New England Biolabs) and *E. coli* BL21-AI (Invitrogen) were used as the cloning and expression hosts respectively, while pHis17 (refer Addgene plasmid #78201 for vector backbone, obtained from Löwe lab, MRC LMB, Cambridge) and pETphos (43) were the plasmid vectors employed for heterologous and inducible overexpression. Cells were grown at 37°C in LB medium containing ampicillin at a final concentration of 100 µg/ml.

For cloning in pHis17, the full-length *frzCD* or DNA coding for *frzCD* variants were amplified with primers designed to ensure insertion of *Nde*1 and *BamH*1 restriction sites at flanking ends of the DNA fragment along the sequence coding for a hexa-histidine tag at the C terminus. The primers as listed in Supplementary Table S3 were used for domain deletions using a restriction free cloning strategy and site-directed mutagenesis (44). For cloning in pETphos, *frzCD* variants were amplified and cloned into pETphos previously digested with NdeI and BamHI. Thus, we obtained ΔMA, ΔH1, ΔH2, ΔH1H2 and FrzCD* (a mutant where three lysines K9, K13 and K18, and two arginines R15 and R17 in the DB region were substituted into glutamates) constructs. Positive clones were confirmed by sequencing and the plasmids were transformed in *E. coli* expression strains such as BL21-AI, BL21-DE3, C43 (BL21-DE3 derivative) cells for checking over-expression.

### Motility phenotypes

For motility phenotypic assays, exponentially growing cells in CYE medium at 32°C were adjusted to an OD_600_ of 10 in TPM buffer and spotted (5 μL) on CYE plates containing an agar concentration of 0.5% or on CF plates containing an agar concentration of 1.5%. The plates without or with vanillate concentration ranging from 0.5 to 50 µM, were incubated at 32°C, and photographed after 48 hours (for twitching motility) or 72 hours (for fruiting body formation) with an Olympus SZ61 binocular.

### Reversal frequencies

Two μl of cells from 4 x 10^8^ cfu.ml^-1^ vegetative CYE cultures were spotted on a thin fresh TPM agar supplied or not with 0.15% IAA. Time-lapse movies were shot for 1 hour with frames captured every 30 seconds. Movies were obtained and reversal frequencies manually counted. Reversal frequencies were measured from cells issued from two biological replicates. Box plots were generated by R and statistical significance was obtained with a Wilcox Test from the R software (https://www.r-project.org/).

### Fluorescence microscopy and image analysis

For fluorescence microscopy analyses, 5 μl of cells from 4 x 10^8^ cfu.ml^-1^ vegetative CYE cultures were spotted on a thin fresh TPM agar pad at the top of a slide. A cover slip was added immediately on the top of the pad, and the obtained slide was analyzed by microscopy using a Nikon TE PFS inverted epifluorescence microscope (100x oil objective NA 1.3 Phase Contrast). To study the colocalization with DNA, the TPM agar pads were supplied with 1 μg/ml DAPI stain and 50 mM vanillate. Prior to imaging, *M. xanthus* cells were grown in 50 mM vanillate. Cell fluorescence profiles were obtained with the “plot profile” function of FIJI. FrzCD cluster position and intensity was represented as demographs and histograms automatically generated with the “MicrobeJ” Fiji/ImageJ plugin created by A. Ducret, Brun Lab (http://www.indiana.edu/~microbej/index.html). All data plots and statistical tests were obtained with the R software (https://www.r-project.org/). Results are issued from biological triplicates.

### Western and methylation analysis

Total cell lysates were analyzed with SDS-PAGE (sodium dodecyl sulfate-polyacrylamide gel electrophoresis) and transferred to a nitrocellulose membrane. After transfer, the membrane was blocked for 2 h at room temperature in 5% powdered milk. Proteins were detected using αFrzCD (1:10,000) (18) from rabbit in a mixture with TBST (Tris-buffered saline with Tween 20; OriGene Technologies Inc., Rockville, MD, USA) in combination with a secondary anti-rabbit antibody (from goat) coupled to HRP (horseradish peroxidase) (1:5,000) (Thermo Fisher Scientific, MA, USA). Methylation was measured by FIJI as percentage of the intensity of the upper band over the intensity of all bands for each given sample. Demethylation was extrapolated as the total fluorescence minus the intensity from the methylated band.

### Protein purification

Proteins expressed from pHis17 plasmid were purified by affinity chromatography (HisTrap, GE Life Sciences) followed by an ion-exchange chromatography step. The temperature was maintained as 4°C throughout. *E. coli* protein expression strains such as BL21-AI, BL21-DE3, C43 (BL21-DE3 derivative) were standardized for each construct and grown in LB broth supplemented with 100 µg/ml ampicillin and incubated at 37°C till the OD_600_ reached 0.6 (mid-exponential phase). The cells were then induced by 0.2% arabinose or 0.5 mM IPTG and incubated at 30°C and 18°C respectively for 5h and 10h. Cells were harvested and resuspended in lysis buffer (200 mM NaCl, 50 mM Tris pH 8 and 10% glycerol) followed by sonication (Sonics VibraCell) at 60% amplitude, pulse 1’’ ON and 3” OFF cycle for 2 minutes) thrice, with an interval of 5 min. The lysate was spun at 35,000 x g for 45 min at 4°C and supernatant was loaded onto a 5-ml HisTrap^TM^ FF pre-equilibrated with buffer A200 (200 mM NaCl, 50 mM Tris pH 8.0). The column was washed extensively with buffer A200, followed by buffer B200 (200 mM NaCl, 500 mM Imidazole, 50 mM Tris pH 8.0), 2% and 5% washes. Next a step gradient of buffer B200, that is, 10%, 20%, 50% and 100% (mixed with A200) facilitated elution of the protein of interest and 1 mM EDTA was added to the fractions to prevent protein degradation. Aliquots were subjected to SDS-PAGE and the fractions containing pure protein were pooled, dialyzed against A50 (50 mM NaCl, 1 mM EDTA, 50 mM Tris pH 8.0). Further, ion-exchange chromatography using MonoQ/MonoS 10/100 GL (GE Healthcare) was performed to remove impurities such as bound DNA. Protein was loaded with A50 buffer and eluted by a gradient of increasing salt concentration ranging from 50 mM to 335 mM (0 to 30 % of A1000 (1000 mM NaCl, 1 mM EDTA, 50 mM Tris pH 8.0) over 20 column volumes). Fractions containing protein were concentrated using centricons (Vivaspin® 10 kDa MWCO or 3 kDa MWCO). Concentrated protein was aliquoted, flash-frozen and stored at -80 °C for further use in experiments shown in Figures 4A, 5 and Supplementary Figure S6, S7. These purified proteins are shown in Supplementary Figure S5A. Proteins expressed from pETphos were purified by affinity chromatography (HisTrap, GE Life Sciences). For protein purification, strains for each construct were grown in LB broth supplemented with 100 µg/ml ampicillin and incubated at 37°C till the OD_600_ reached 0.6. The cells were then induced by 0.5 mM IPTG and incubated at 18°C for 10h. Cells were harvested and resuspended in lysis buffer (300 mM NaCl, 50 mM Tris pH 7.4) followed by two French press cycles. The lysate was spun at 18,000 x g for 20 min at 4°C and supernatant was loaded on to a 5-ml column containing 1 ml of agarose beads pre-equilibrated with Ni^2+^ and lysis buffer. The column was washed extensively with lysis buffer containing 10 mM and then 75 mM imidazole. Elution was carried with lysis buffer containing 200 mM or 500 mM imidazole. Aliquots were subjected to SDS-PAGE and the fractions containing pure protein were pooled, dialyzed against a buffer containing 150 mM NaCl, 10 mM Tris pH 7.4. Fractions containing protein were concentrated using centricons (Vivaspin® 3 kDa MWCO). Concentrated protein was aliquoted, flash frozen and stored at -80 °C for further use in experiments shown in Figures 4B. These proteins are shown in Supplementary Figure S5B.

### Circular dichroism spectroscopy

Far-UV CD measurements were carried out on a Jasco J-720 spectropolarimeter. The parameters used for measurements of far-UV CD spectra were as follows: step resolution, 1 nm; scan speed, 100 nm/min; and bandwidth, 1 nm. The concentration of FrzCD and DMA were 2.5 µM and 20 µM respectively. CAPITO was used to analyze and plot CD data (45).

### Electrophoretic mobility shift assay (EMSA)

EMSAs were carried out for varying lengths of DNA including 8, 12, 20, 35, 69, 432 and 600 bp of which, 432 and 600 bp were PCR amplicons while the remaining DNA fragments were custom-synthesized (Sigma and Eurofins) and annealed. A single stranded (ss) poly A consisting of 8 bases was also used and all DNA sequences used for the study have been listed in Table S4. Increasing concentrations of purified proteins up to 50 µM were incubated with 0.5 to 2 µM of DNA at 25°C for 30 min in a reaction buffer comprising 50 mM NaCl, 1 mM dithiothreitol, 10% glycerol and 50 mM Tris pH 7.4. A higher concentration of DNA, 2 µM, was used for 8-bp DNA to allow visualization on agarose gel. The samples were mixed with gel loading buffer and loaded on to the appropriate percentage agarose gel containing ethidium bromide. For instance, 4 % agarose gels were used for 8-bp, 12-bp, 20-bp and 35-bp DNA, while 2 % were used for 69-bp and 432-bp DNA. Samples with short DNA fragments were electrophoresed at 80-90 V for 30 min in 1 x TAE (40 mM Tris-acetate, 1 mM EDTA) buffer, followed by visualization under UV light. For longer DNAs, the gels were run at 100 V from 45 min to 1 hr. The gels were observed for a higher shift in the DNA bands in presence of increasing protein concentrations.

### Biolayer interferometry

Protein-DNA interaction experiments were conducted at 25°C with the BLItz instrument from ForteBio (Menlo Park, CA, USA) as described previously (19, 24). A 178 bp DNA fragment was amplified with a forward primer conjugated at the 5’ with biotin (Eurogentec) and immobilized onto a streptavidin biosensor (ForteBio) in duplicate at a 5 nM concentration. Purified proteins were used as the analyte throughout the study at concentrations from 0.7 to 17 mM. The assay was conducted in 10 mM Tris pH 7.4, 150 mM NaCl. The binding reactions were performed with an initial baseline during 30 sec, an association step at 120 sec and a dissociation step of 120 sec, with lateral shaking at 2,200 rpm. A sensor reference subtraction was applied to account for non-specific binding, background, and signal drift to minimize sensor variability.

### SEC-MALS (Size Exclusion Chromatography with Multi Angle Light Scattering)

SEC coupled with multi angle light scattering was employed to estimate the molar mass of FrzCD proteins in solution. Superdex 200 Increase 10/300 GL (GE Healthcare) column coupled to Wyatt Dawn HELIOS II equipped with light scattering detectors at 18-angles along with a differential refractive index detector (Optilab TrEX) was used for the analysis. The column was equilibrated with buffer containing 50 mM NaCl and 50 mM Tris pH 8.0 at room temperature (approximately 25°C), followed by injection of 100 µl of 45 µM and 150 µM (FrzCD); 70 µM and 200 µM (DMA); 25 µM (DH1H2); 25 µM and 65 µM (DH2); 25 µM (DH1). All the runs were carried out at a flow rate of 0.4 ml/min. The observed molecular masses for proteins were estimated at various points along the peaks in elution curves, based on the differential refractive index and scattered intensities. Zimm model of ASTRA software available with the equipment was used for fitting the data. Observed molecular masses were compared with the theoretical masses to estimate oligomeric status of each protein. Bovine serum albumin (2 mg/ml) was used for calibration of the system.

Further, to determine a change in the oligomeric status of proteins upon DNA binding, runs were carried out with the ligand, that is, 35-bp DNA was included in the injected samples. Hence, 100 µl of 70 µM FrzCD with 30 µM DNA; 200 µM FrzCD with 100 µM DNA; 200 µM DB-H1-H2 with 100 µM DNA; 100 µM DB-H1-MA with 50 µM DNA were injected while 30 µM DNA and 70 µM FrzCD served as control runs. Observed molecular masses for all the peaks in elution curve were estimated and compared with theoretical masses of probable protein-DNA species.

## Acknowledgments

We thank Pravin Dewangan, Sangita Niranjan and Radha Chauhan, NCCS Pune for use of the SEC-MALS facility, Sandhya Bhatia and Jayant Udgaonkar for access to CD spectroscopy facility, Somya Madan for conducting initial trials of FrzCD modeling, and Kaustubh Amritkar and Suman Pal for preliminary EMSA experiments. We also thank Deborah Byrne from the Protein Expression Platform (CNRS, IMM, Marseille) for greatly helping with the BLItz experiments and Yann Denis from the Transcriptomic Platform (CNRS, IMM, Marseille) for analyzing the FrzE expression levels. We thank Vladimir Pelicic and Julien Herrou for their critical reading of the manuscript.

