## Supplementary material for "Di-HAMP domains of a cytoplasmic chemoreceptor modulate nucleoid array formation and downstream signalling"

**This PDF file includes:**

Supporting text

Figures S1 to S7

Tables S1 to S5

**Supplementary Figures**

**Figure S1**

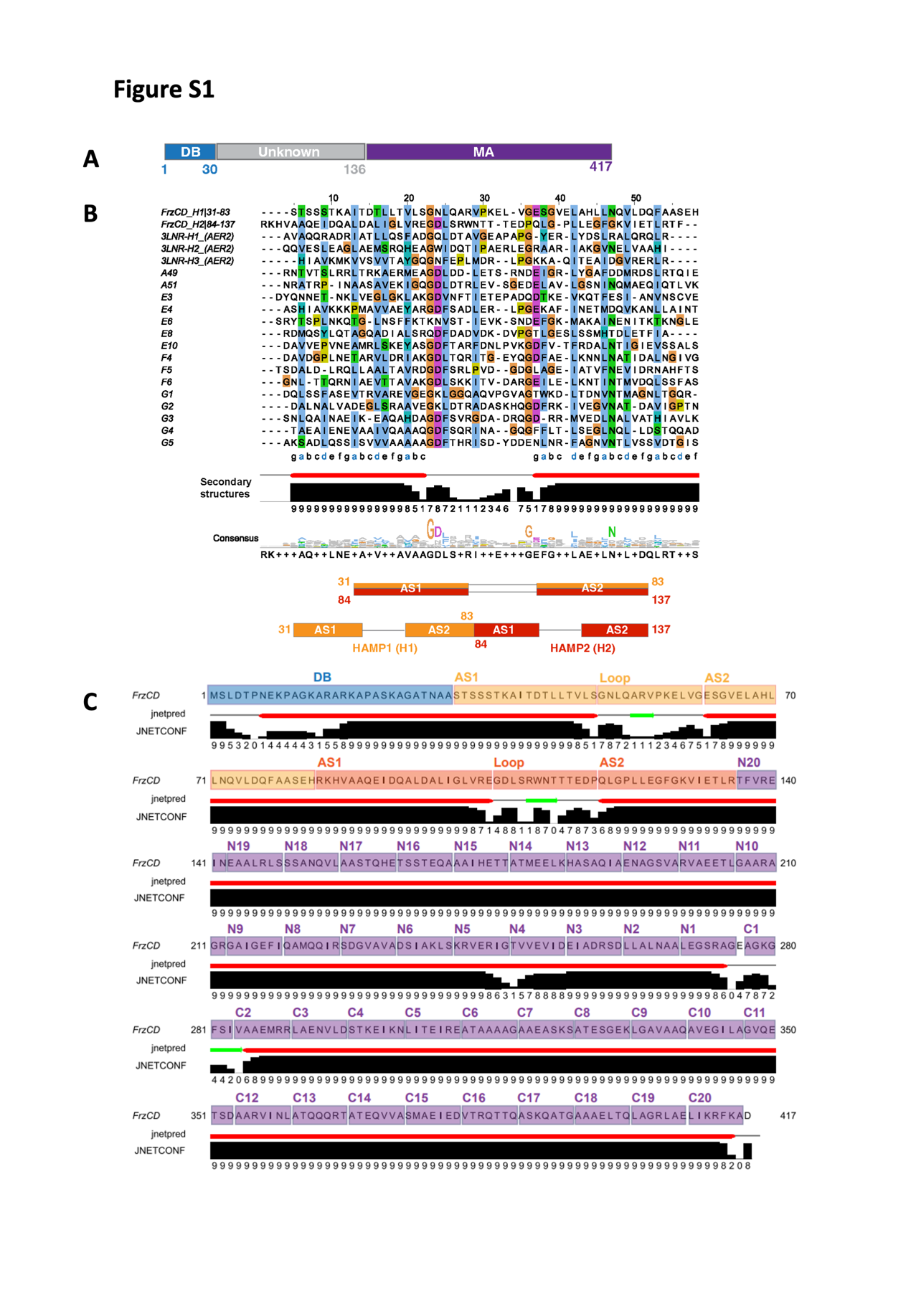

**Supplementary Figure S1: FrzCD is mostly composed of alpha helices and its N-terminal domain comprises tandem HAMP domains.**

**A.** FrzCD comprises an N-terminal DB domain (1-30) and a C-terminal MA domain (137-417). Residues 31 to 137 are of unknown fold as inferred from sequence-based predictions.

**B.** Sequence alignment (top) and secondary structure (bottom) suggest the presence of two concatenated HAMP domains in FrzCD (H1: residues 31-83 and H2: residues 84-137). The aligned sequences are the consensus sequences of various classes of HAMP domains. Color scheme as per ClustalX is used for representation in the alignments. Hydrophobic residues of helices AS1 and AS2 of H1 and H2 are labeled in blue in the alignment. The conserved glycines flanking the loop region are in orange. The AS2 of H1 is in tandem with AS1 of H2. The confidence of secondary structure prediction is shown by numbers ranging from 0 to 9 (lowest to highest).

**C.** Secondary structure prediction of FrzCD. DB (blue), H1 and H2 (each consisting of AS1, loop and AS2; orange and red), and MA (20 heptads of the N and C-terminal ends labeled as N1-20 and C1-20; purple) domains are marked on the amino acid sequence. Helices and strands are shown as red cylinders and green arrows, respectively. The confidence of secondary structure prediction is shown by numbers ranging from 0 to 9 (lowest to highest).

**Figure S2**
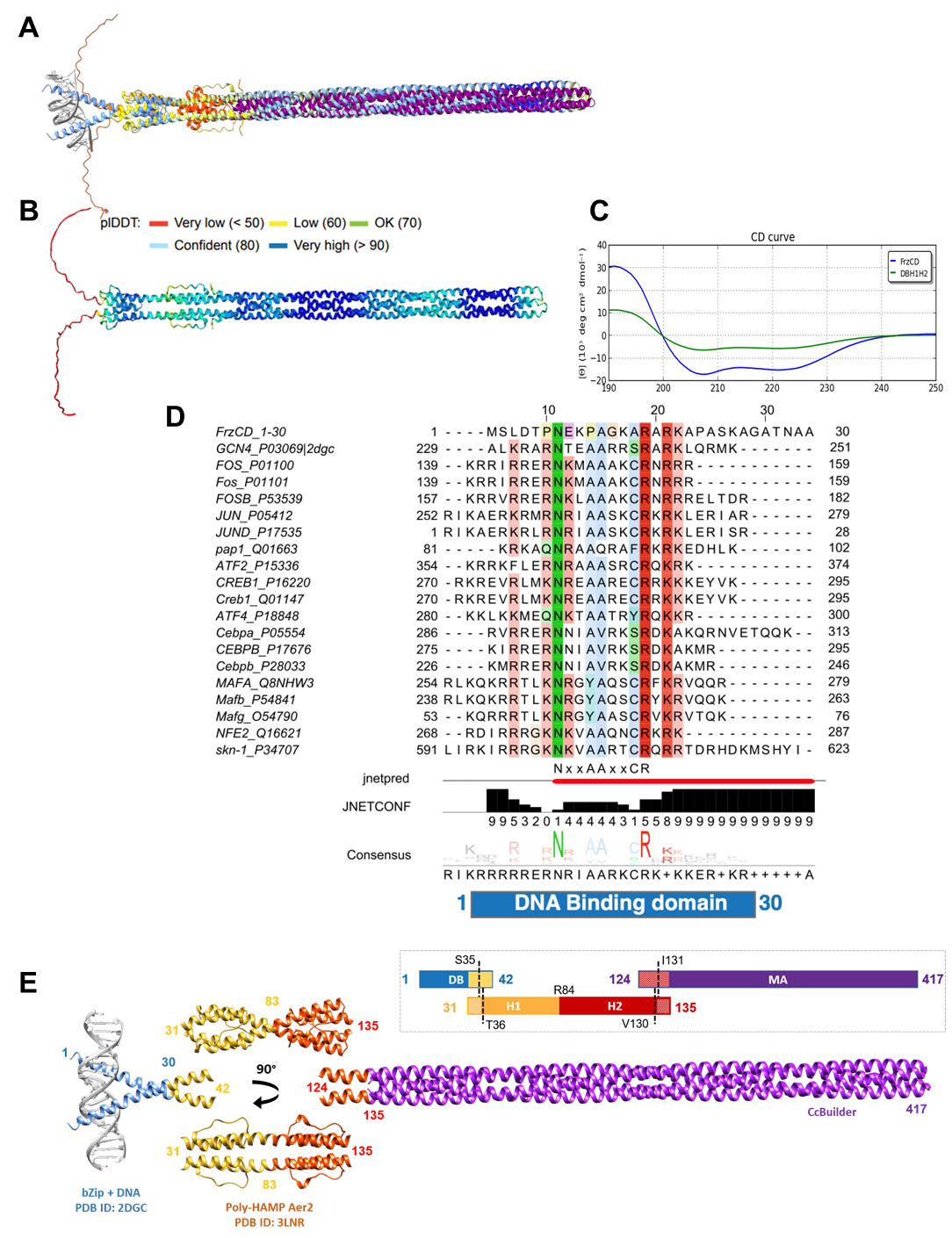

**Supplementary figure S2. Validation of homology model of FrzCD**.

**A.** Superposition of the AlphaFold coordinates of FrzCD dimer mode (color-coded according to reliability scored by plDDT) with our homology model (color-coded according to domains; DB: blue, H1: golden, H2: red, MA: purple)

**B**. FrzCD AlphaFold model color-coded according to reliability scored by plDDT.

**C.** Circular dichroism spectroscopy of FrzCD (blue) and DB-H1-H2 (green) shows presence of extended helical regions (71% and 44% helical content respectively).

**D.** DNA binding domain of FrzCD is similar to bZIP DNA binding sequences. Sequence alignment of the DB domain with leucine zipper domains highlights the conserved consensus sequence NxxAAxxCR (where x is any residue) and positively charged residues (R and K). Secondary structure prediction of the DB domain highlights a helical conformation similar to basic leucine zipper proteins. Color scheme as per ClustalX is used for representation in the alignments.

**E.** Schematic representation of the strategy employed to generate a homology model of FrzCD where individual domains were modeled separately with an overlap of 12 amino acids for ease of stitching. The C-⍺ of the 12 overlapping residues between each of the models were superimposed followed by connecting S35 and T36 of DB and H1, and V130 and I131 of H2 and MA, respectively, and deleting the overlapping residue coordinates. Homology models for the DNA binding and di-HAMP domains were generated using leucine zipper (PDB ID: 2DGC) and Aer2 (PDB ID: 3LNR) respectively, in SWISS-MODEL while the MA domain was modeled using CCBuilder. Model is color-coded according to domains (DB: blue, H1: golden, H2: red, MA: purple).

**Figure S3**
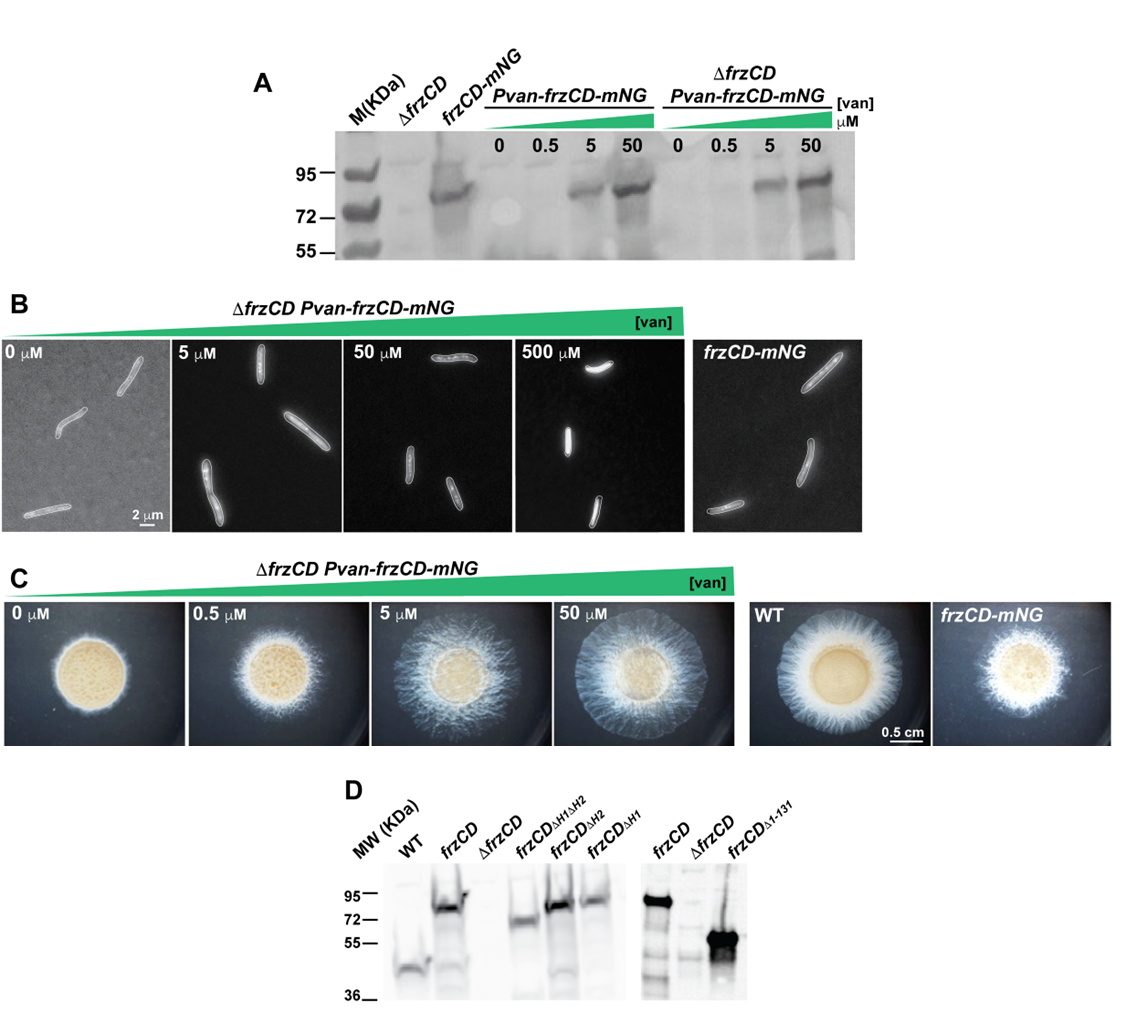

**Supplementary Figure S3. FrzCD-mNG forms clusters at the nucleoid.**

**A.** Representative Western blot with α-FrzCD antibodies on whole cell extracts of strains DZ2 (wild type), EM525 (*ΔfrzCD*), EM724 (*frzCDmNG*) and EM885 (*ΔfrzCD* *pvan-frzCDmNG*). *pvan-frzCDmNG* and *ΔfrzCD* *pvan-frzCDmNG* were grown in the presence of the indicated concentrations of vanillate.

**B.** *ΔfrzCD* *pvan-frzCDmNG* was grown in the presence of the indicated concentrations of vanillate and imaged using a fluorescence microscope.

**C.** The same cells were used for motility phenotypes on 0.5% CYE agar and imaged at 48h.

**D.** Western blot with α-FrzCD antibodies on cell extract of strains DZ2 (wild type), EM525 (*ΔfrzCD*), EM885 (*ΔfrzCD pvan-frzCDmNG*), EM914 (*ΔfrzCD pvan-frzCDmNG^ΔH1^*), EM913 (*ΔfrzCD pvan-frzCDmNG^ΔH2^*), EM911 (*ΔfrzCD pvan-frzCDmNG^ΔH1ΔH2^*) and EM908 (*ΔfrzCD pvan-frzCDmNG^Δ1-131^*).

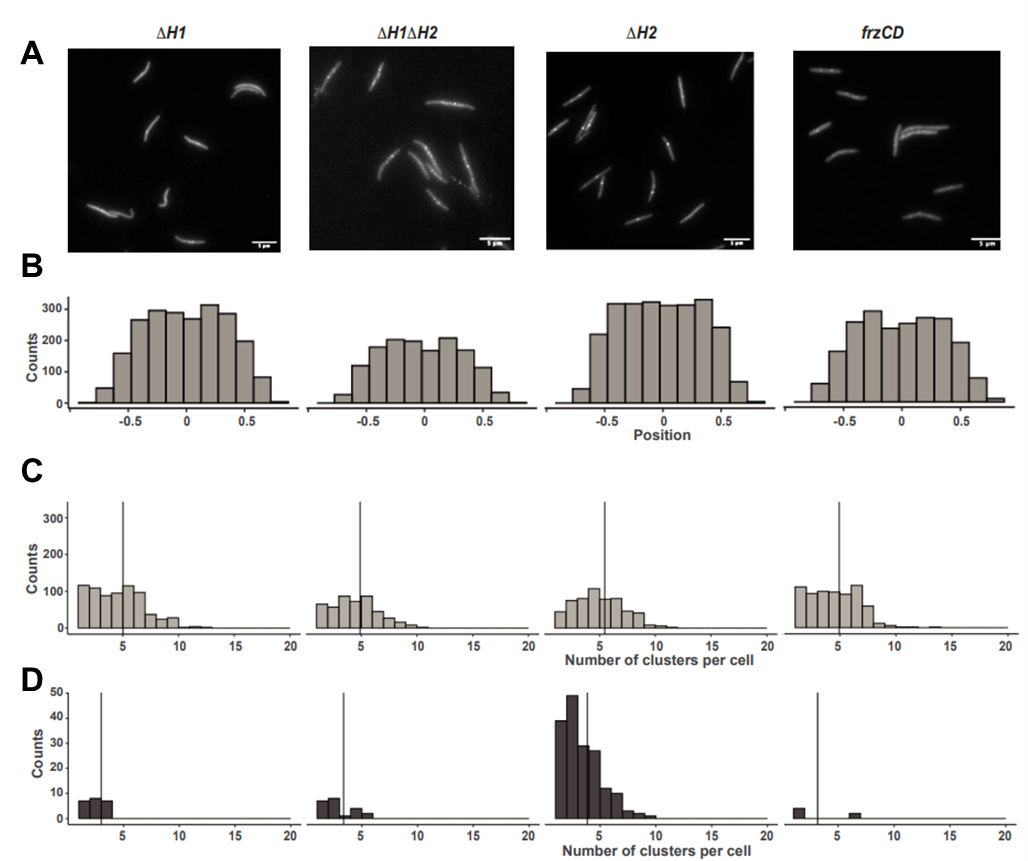
**Figure S4**

**Supplementary Figure S4. Deletions of different HAMP domains have different effects on clusters number and intensity.**

**A.** Large views of micrographs of *M. xanthus* cells from strains EM885 (*ΔfrzCD pvan-frzCDmNG*), EM914 (*ΔfrzCD pvan-frzCDmNG^ΔH1^*), EM913 (*ΔfrzCD pvan-frzCDmNG^ΔH2^*) or EM911 (*ΔfrzCD pvan-frzCDmNG^ΔH1ΔH2^*). Scale bars correspond to 5 μm.

**B.** Absolute number of clusters in function of their position in cells (from -1 to +1 on the y axis). “0” is the cell center; 0.5 and -0.5 are quarter positions.

**C.** Absolute number of low-fluorescence clusters per cell in function of their fluorescence intensity (arbitrary units).

**D.** Absolute number of high-fluorescence clusters per cell in function of their fluorescence intensity (arbitrary units).

**Figure S5**

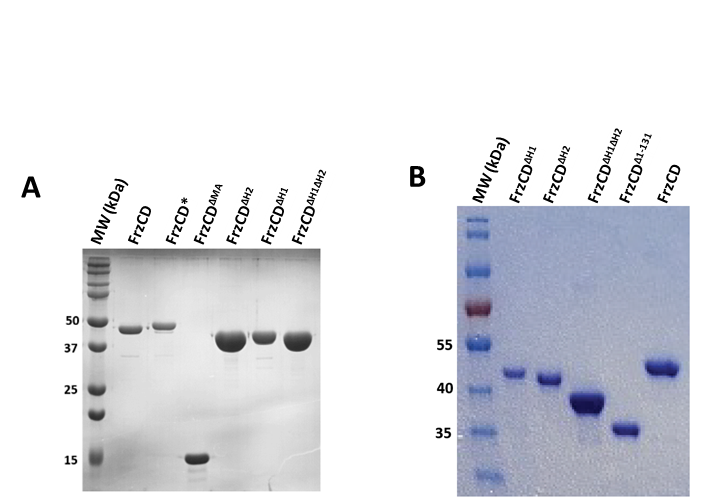

**Supplementary Figure S5. The proteins used in this study are stably expressed.**

**A, B.** 12 % **(A)** or 10 % **(B)** SDS-PAGE profiles of the purified FrzCD constructs used in this study. Proteins in (A) were used to run the experiments shown on Figures 4, Figure 5, and Supplementary Figure S6. Proteins in (B) were used to run the experiments shown on Figures 4 (B-D).

**Figure S6**

**Supplementary Figure S6. The MA domain stabilizes FrzCD binding to longer DNA fragments.**

**A-B.** Representative electrophoretic mobility shift assays (EMSA) resolving the FrzCD and FrzCD^ΔMA^ binding to DNA. The protein concentrations of purified FrzCD protein domains are indicated corresponding to each lane and were incubated with the indicated concentration and length of DNA fragments from 8 to 432 bp.

**Figure S7**
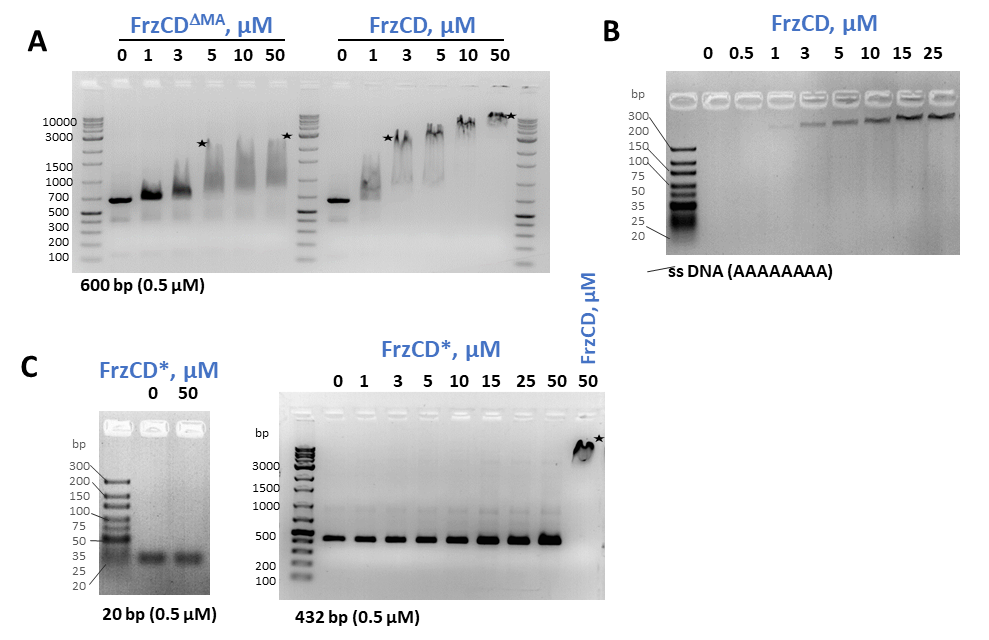

**Supplementary Figure S7. Characteristics of FrzCD DNA binding**

**A.** MA domain influences stable supershifts of longer DNA (binding profile comparison of 600-bp is shown).

**B.** FrzCD binds to single-stranded DNA. 1 µM of 8-base poly-A ssDNA titrated with increasing FrzCD concentration from 0 to 25 µM, according to lane labels.

**C.** FrzCD* shows no binding to DNA, and hence no shift is observed in EMSA.

**Supplementary Tables**

**Supplementary Table S1. Strains used in this study**

| ***M. xanthus*** | **Genotype** | **Reference or Source** |
| --- | --- | --- |
| EM814 | *DZ2*, Wild type | Campos and Zusman, 1975 |
| EM525 | *ΔfrzCD* | Bustamante et al., 2004 |
| EM527 | *ΔfrzF* | Bustamante et al., 2004 |
| EM528 | *ΔfrzG* | Bustamante et al., 2004 |
| EM724 | *frzCD-mNeongreen* | This study |
| EM777 | *frzCD^ΔH1^-mNeongreen* | This study |
| EM778 | *frzCD^ΔH2^-mNeongreen* | This study |
| EM779 | *frzCD ^ΔH1H2^-mNeongreen* | This study |
| EM889 | *frzCD^ΔH1^* | This study |
| EM775 | *frzCD^ΔH2^* | This study |
| EM776 | *frzCD ^ΔH1H2^* | This study |
| EM886 | *P^van^-frzCD-mNeongreen* | This study |
| EM885 | *ΔfrzCD, P^van^-frzCD-mNeongreen* | This study |
| EM914 | *ΔfrzCD, P^van^-frzCD ^ΔH1^-mNeongreen* | This study |
| EM913 | *ΔfrzCD, P^van^-frzCD ^ΔH2^-mNeongreen* | This study |
| EM911 | *ΔfrzCD, P^van^-frzCD ^ΔH1H2^-mNeongreen* | This study |
| ***E. coli*** | **Genotype** | **Reference or Source** |
| DH5a | *F- Φ80lacZΔM15 Δ(lacZYA-argF) U169 recA1 endA1 hsdR17 (rK-, mK+) phoA supE44 λ- thi-1 gyrA96 relA1* | New England Biolabs |
| BL21-AI | F^-^*omp*T *hsd*S_B_ (r_B_^-^ m_B_^-^) *gal dcm ara*B::T7RNAP-*tet*A | Invitrogen |

**Supplementary Table S2. Plasmids used in this study**

| **Plasmid** | **Insert** | **Source** |
| --- | --- | --- |
| pKY480 | Empty vector | Mauriello et al., 2009 |
| pEM517 | pKY480 with *frzCD-mNeongreen* | This study |
| pBJ114 | Empty vector | Bustamante et al., 2004 |
| pEM566 | pBJ114 with *frzCD^ΔH1^* | This study |
| pEM567 | pBJ114 with *frzCD^ΔH2^* | This study |
| pEM569 | pBJ114 with *frzCD^ΔH1 ΔH2^* | This study |
| pMR3690 | Empty vector | Iniesta et al., 2012 |
| pEM627 | pMR3690 with *frzCD-mNeongreen* | This study |
| pEM646 | pMR3690 with *frzCD^ΔH1^-mNeongreen* | This study |
| pEM643 | pMR3690 with *frzCD^ΔH2^-mNeongreen* | This study |
| pEM644 | pMR3690 with *frzCD^ΔH1H2^-mNeongreen* | This study |
| FrzCD | pHis17 with *frzCD* | This study |
| FrzCD^ΔMA^ | pHis17 with *frzCD^ΔMA^* | This study |
| FrzCD^ΔH1^ | pHis17 with *frzCD^ΔH1^* | This study |
| FrzCD^ΔH2^ | pHis17 with *frzCD^ΔH2^* | This study |
| FrzCD^ΔH1H2^ | pHis17 with *frzCD^ΔH1H2^* | This study |
| FrzCD* | pHis17 with *frzCD^K9E_K13E_R15E_R17E_K18E^* | This study |
| pETphos | Empty vector | Canova et al., 2008 |
| pEM410 | pETphos with *frzCD* | Moine et al., 2017 |
| pEM414 | pETphos with *frzCD^Δ1-131^* | Moine et al., 2017 |
| pEM663 | pETphos with *frzCD^ΔH1^* | This study |
| pEM662 | pETphos with *frzCD^ΔH2^* | This study |
| pEM658 | pETphos with *frzCD^ΔH1H2^* | This study |

**Supplementary Table S3. Primers used for restriction-free cloning of domain-wise deletion constructs**

| **Primer** | **Sequence (5’ to 3’)** |
| --- | --- |
| FrzCD^K9E_K13E_R15E_R17E_K18E^_f | CCCCCAACGAGGAGCCCGCTGGCGAGGCTGAAGCCGAGGAGGCCCCCGCCTCCGAGGCCGCGGCC |
| FrzCD^ΔH1^_f | GGCGCCACGAACGCGGCGTCGCGCAAGCATGTGGCGGCG |
| FrzCD ^ΔH1ΔH2^_f | GGCGCCACGAACGCGGCGTCGGTGCGGGAGATCAACGAG |
| FrzCD^ΔH2^_f | CAGTTCGCGGCCTCCGAGCACGTGCGGGAGATCAACGAG |
| FrzCD^ΔMA^_r | GCTTTTAATGATGATGATGATGATGGGATCCGAAGGTGCGCAGCGTCTCGATG |

**Supplementary Table S4. List of DNA sequences used for EMSA**

| **DNA length** | **Sequence (5’ to 3’)** |
| --- | --- |
| 8bp | ACTGCAGT |
| ss DNA | AAAAAAAA |
| 12 bp | CTCACTATAGGG |
| 20 bp | TAATACGACTCACTATAGGG |
| 35 bp | GTCACCTGCTCTAGCTAATAGACTGAGCCGAGGTG |
| 69 bp | CTTGCAGTAGAGCTGACCATGATTACGCCATCAGCAGCTCCAGGTCGTACCTCCAGCTACCAATCCCCG |
| 178bp | GGATCGCATCTGCCTACATCCCCAGCTCCCGGTCGGTCCGCTCGGAACCTAGCCCGGGTCAAAGACCCGGGGTCTATGTTGACCCATTTGGTGGGGATGGGTCTAATTGGACCTGGCGTTTTTTCGCGCCATGGAAATGTCAAGGCCCGTGATTCCAGACTGTTGGGCAGGGAGTT |
| 432 bp | TAATACGACTCACTATAGGGAGACCACAACGGTTTCCCTCTAGAAATAATTTTGTTTAACTTTAAGAAGGAGATATACATATGTCCCTGGACACCCCCAACGAGAAGCCCGCTGGCAAGGCTCGCGCCCGGAAGGCCCCCGCCTCCAAGGCCGGCGCCACGAACGCGGCGTCGACCTCTTCCTCCACCAAGGCCATCACCGACACGCTGCTGACGGTGCTGTCCGGCAACCTGCAGGCCCGCGTGCCCAAGGAGCTGGTCGGTGAGTCCGGCGTGGAGCTGGCGCACCTGCTCAACCAGGTGCTGGACCAGTTCGCGGCCTCCGAGCACCGCAAGCATGTGGCGGCGCAGGAGATCGACCAGGCGTTGGATGCGCTCATCGGCCTGGTGCGCGAGGGCGGATCCCATCATCATCATCATCATTAAAAGC |

**Supplementary Table S5: Summary of the SEC-MALS profile peaks and their corresponding expected and estimated molecular masses.**

| **Panel peak number** | **Probable Protein/DNA/Protein-DNA species** | **Theoretical Molar Mass (kDa)** | **Observed Molar Mass (kDa)** |
| --- | --- | --- | --- |
| A | FrzCD dimer (89 kDa) +35-bp DNA (22 kDa) | 111 | 111 ± 0.3 |
|  | FrzCD dimer (89 kDa) | 89 | 89 |
|  | 35-bp DNA (22 kDa) | 22 | 22 |
| B | FrzCD^ΔMA^ (31 kDa) + 35-bp DNA (22 kDa) | 53 | 47 ± 2 |
| C.1 | 35-bp DNA (22 kDa) | 22 | 21.8 ± 0.3 |
| C.2 | FrzCD dimer (89 kDa) | 89 | 85.8 ± 1.2 |
| C.3 | Higher order species |  | 138.2 ± 1.2 |
| C.4 | Higher order species |  | 199 ± 5 |
| D.1 | 35-bp DNA (22 kDa) | 22 | 24.7 ± 0.7 |
| D.2 | FrzCD^ΔH2^ (80 kDa) + 35-bp DNA (22 kDa) | 102 | 107 ± 5.2 |
| D.3 | Higher order species (broad peak) |  | 194 ± 12 |
